# Long-term realized genetic gain and population dynamics under genomic selection in Brazilian cassava Germplasm

**DOI:** 10.64898/2026.07.18.739356

**Authors:** Gabriel Mamedio de Freitas, Diana C. Solarte Certuche, Jean-Luc Jannink, Eder Jorge de Oliveira, Antonio Augusto Franco Garcia

## Abstract

Genomic selection has become an important strategy in cassava breeding, enabling faster selection cycles and sustained genetic progress. Despite its widespread adoption, long-term evaluations integrating predictive performance, realized genetic gain, and genetic diversity remain scarce, particularly in clonally propagated crops. We present a comprehensive assessment of genomic selection outcomes in the Brazilian cassava breeding program across four recurrent selection cycles (C0 to C3) implemented between 2011 and 2024, using historical phenotypic and genomic data from 210 multi-environment trials. Predictive ability of genomic best linear unbiased prediction models ranged from low to moderate, depending on the trait’s genetic architecture and heritability. Prediction accuracies were highest in early cycles (C0 and C1) and showed modest declines in later cycles (C2 and C3). Root yield, shoot yield, plant height, starch content, and dry matter content exhibited stable predictive performance across cycles, with a gradual reduction in RMSE, indicating improved model calibration as training populations expanded. Regression analyses of genomic estimated breeding values revealed significant realized genetic gains for most yield-related traits. In contrast, dry matter content and starch content exhibited small, non-significant negative trends, consistent with known unfavorable genetic correlations with yield. Targeted reductions in plant architecture scores reflected deliberate selection for ideotypes suited to mechanized production systems. At the same time, analyses of genetic diversity revealed a slight decrease in observed heterozygosity, with higher values in the most advanced selection cycle. These results provide an integrated framework for monitoring predictive performance, realized genetic gain, and population genetic dynamics under long-term genomic selection. Collectively, they offer valuable insights into balancing short-term genetic improvement with long-term sustainability and support the development of strategies to optimize selection decisions, breeding planning, and population management in Brazilian cassava breeding programs.

## Introduction

Cassava (*Manihot esculenta* Crantz) is a crop of South American origin, with its primary center of domestication located in the Brazilian Central-West region, and it subsequently experienced extensive global dissemination across tropical regions from the mid-16th to the 19th centuries (Gleadow, Maher, and Cliff, 2023). Owing to this history of domestication and diffusion, cassava is currently cultivated across a wide range of tropical and sub-Saharan environments. In recent decades, the crop has received increasing scientific and strategic attention due to its exceptional ability to produce high amounts of carbohydrates under low-input conditions, particularly in marginal environments where other staple crops perform poorly (Fathima et al., 2023; Wang et al., 2014). Moreover, cassava exhibits notable resilience to biotic stresses, including major pests and diseases, as well as tolerance to adverse conditions associated with climate variability and change. These attributes position cassava as a key crop for food security (Gleadow, Maher, and Cliff, 2023; Ceballos, Iglesias, et al., 2004) and as an important raw material for the industrial sector in Brazil (Andrade, Sousa, Oliveira, et al., 2019b; Oliveira, Resende, et al., 2012).

Despite its agronomic advantages, cassava also presents substantial challenges for genetic improvement. The species is characterized by high levels of heterozygosity, a long cultivation cycle, difficulties in controlled hybridization due to floral biology constraints, and limited availability of vegetative planting material. Together, these factors negatively affect experimental precision and slow the generation of reliable phenotypic data in field trials. As a consequence, under conventional breeding schemes, cassava improvement cycles are prolonged and may span 8 to 10 years from crossing to cultivar release (Ceballos, Iglesias, et al., 2004; Ceballos, Rojanaridpiched, et al., 2020). In response to these limitations, Embrapa has invested sustained efforts over the past 15 years to modernize cassava breeding pipelines, combining classical breeding strategies (Nascimento, Andrade, et al., 2024; Sampaio Filho et al., 2023a; Oliveira, Oliveira, and Boas, 2017) with the progressive incorporation of genomic selection (GS) approaches (Oliveira, Resende, et al., 2012; Torres et al., 2019; Andrade, Sousa, Wolfe, et al., 2022; Andrade, Sousa, Oliveira, et al., 2019a; Costa et al., 2024).

GS relies on genome-wide molecular marker information to estimate breeding values for genotyped and phenotyped individuals, as well as to predict the genetic merit of individuals that have been genotyped but not yet phenotyped (Meuwissen, Hayes, and Goddard, 2001). Since its conceptualization, GS has expanded the scope of predictive breeding and has been progressively integrated into plant breeding programs worldwide (Campos et al., 2017; Crossa et al., 2017), becoming a routine component of modern breeding strategies. In the Embrapa cassava breeding program, GS was first implemented in 2011 (Oliveira, Resende, et al., 2012), resulting in more than 13 years of continuous research and the establishment of four recurrent genomic selection populations, the most recent of which is still under field evaluation.

In this context, genomic selection models enable prediction of genetic values that may incorporate both additive and dominance effects, depending on the model’s structure and assumptions. These predictions can be used directly for selection or integrated into selection indices, with specific weights assigned to different traits according to the breeding objectives. This approach enables prioritizing attributes associated with the development of the desired end product while maximizing expected genetic gain for the target traits within the adopted selection framework (Andrade, Sousa, Oliveira, et al., 2019a; Andrade, Sousa, Wolfe, et al., 2022; Costa et al., 2024).

However, as breeding programs advance through successive selection cycles, their objectives often evolve in response to market demands, production systems, and end-user preferences. Consequently, breeding efforts increasingly aim to deliver products that simultaneously meet the criteria of market acceptability and agronomic adaptability, using metrics relevant to both consumers and producers (Covarrubias-Pazaran, 2020; Seck et al., 2023). In this context, quality indicators can be defined to evaluate the reliability and efficiency of the breeding pipeline relative to its objectives, either within a single cycle (through predicted gains) or across multiple selection cycles. For the latter purpose, realized genetic gain (RGG) is a particularly informative metric, as it quantifies the cumulative response to selection and monitors the consistency of genetic progress over time (Covarrubias-Pazaran, 2020; Seck et al., 2023; Dieng et al., 2024; Rutkoski, 2019a).

Realized genetic gain is a quantitative measure of the progress achieved through coordinated selection practices in a breeding program. It is commonly estimated via linear regression of trait values or selection indices against time or selection cycles, and may be calculated using the phenotypic best linear unbiased estimator (BLUE) or predictor (BLUP), as well as GEBVs estimated across the evaluated cycles. The regression slope represents the absolute gain per cycle, directly reflecting the cumulative effect of selection over time. When expressed relative to the regression intercept, this slope provides a standardized estimate of genetic gain in percentage terms, enabling meaningful comparisons of genetic progress across cycles (Seck et al., 2023; Rutkoski, 2019b; Dieng et al., 2024).

The incorporation of genomic selection into the national cassava breeding program has also highlighted the need to monitor additional strategic indicators beyond short-term genetic gain. Although GS, combined with high selection intensity, can substantially reduce breeding cycle length (Crossa et al., 2017), sustained genetic progress depends on maintaining adequate genetic diversity. Previous studies, such as Rutkoski et al. (2015), have demonstrated that genomic selection may reduce genetic diversity across breeding cycles if not properly managed. Therefore, diversity metrics, including expected heterozygosity and inbreeding coefficients, should be routinely monitored to prevent genetic erosion that could lead to inbreeding depression and compromise long-term genetic gains in priority traits.

Evaluations of realized genetic gain still present an important gap in cassava breeding, particularly regarding their application in conjunction with genomic selection. Realized genetic gain has previously been assessed by Delgado et al. (2024) and Manze et al. (2021), who analyzed genetic progress across multiple breeding cycles in two independent programs. However, these studies relied exclusively on phenotypic estimates (BLUEs and BLUPs) and did not incorporate genomic estimated breeding values (GEBVs).

The incorporation of information from molecular markers, in addition to enabling indirect selection, extends analyses to more robust inferences about genetic progress over time (Rutkoski, 2019b; Covarrubias-Pazaran, 2020). Moreover, this approach provides a more comprehensive representation of the overall breeding program by enabling the integration of population genetics analyses to evaluate the feasibility and effectiveness of the selection strategies adopted, as well as to identify adjustments and changes that can be implemented based on the efficiency of realized genetic gains for a given trait, including the use of computational simulations as decision-support tools (Rutkoski, 2019a; Seck et al., 2023).

In this context, an integrated assessment of quality indicators is essential to evaluate the consistency, efficiency, and sustainability of breeding efforts conducted over 13 years of genomic selection in the current program, as well as their implications for the genetic diversity of the resulting germplasm. Accordingly, the present study analyzes historical data from 210 clonal evaluation trials conducted under the GS framework between 2011 and 2024, with the following objectives: (i) to assess realized predictive ability and the RMSE of implemented genomic selection cycles using historical data; (ii) to estimate realized genetic gain based on GEBVs for 13 root yield and quality traits; and (iii) to evaluate changes in genetic diversity across the four genomic selection cycles implemented during this period.

## Methods & Materials

### Germplasm and recurrent-selection pipeline overview

The materials used in this study come from the Embrapa cassava improvement program and comprise the historical series of recurrent genomic-selection populations implemented between 2011 and 2024, including 210 clonal evaluation trials and four recurrent selection populations established during this period. The recurrent selection pipeline implemented by Embrapa combines (i) multi-environment clonal phenotyping, (ii) medium-density genotyping of breeding material, and (iii) genomic prediction using both additive and non-additive models.

#### Constitution of the training population, Cycle 0 (C0)

The initial training population (C0) was assembled from Embrapa’s elite germplasm pool, advanced clones, and breeder selections that were evaluated during the program’s early operational years (2011–2013). C0 aggregates the historical phenotypes and genotypes available at program start and was used to train prediction models applied in subsequent cycles. Clonal evaluations that contributed to C0 were conducted across representative agroecological zones for Brazilian cassava, using replicated clonal trials. Experimental designs were predominantly augmented or randomized complete block designs with 2–3 replications per environment; plot size and agronomic management followed Embrapa standard protocols (Dos Santos et al., 2023b; Guimarães et al., 2025; Fukuda et al., 2010).

The traits used to build the training set reflect the study objectives (root yield, root number, dry matter content, and other additional root quality/production traits). Phenotypes were recorded at physiological maturity following standardized harvest protocols; fresh root weight per plot was converted to per-plant estimates when required.

All C0 clones were genotyped using a reduced-representation sequencing approach, such as Genotyping-by-Sequencing (GBS) (Elshire et al., 2011) and/or Diversity Arrays Technology (DArT-seq), depending on program availability and year. Marker filtering removed SNPs with minor allele frequency (MAF) < 0.05, call rate < 80%, and markers with extreme segregation distortion; individuals with ≥ 20% missing genotypes were excluded from model training. Missing genotypes were imputed using the Beagle population-aware algorithm (Browning and Browning, 2016), and a genomic relationship matrix was computed from the filtered marker set for use in genomic models. Quality control and marker-processing scripts were implemented in PLINK and in custom R pipelines, following Purcell et al. (2007). To quantify baseline predictive ability and to guide model selection, C0 was partitioned in multiple cross-validation schemes that mimic operational decision contexts. Predictive ability (correlation between predicted and observed values) and error (RMSE) were the primary metrics used to rank models for operational deployment.

#### Formation of Cycle 1 (C1)

Candidate parents for the first recurrent cycle (C1) were selected from superior clones of C0 using a multi-trait selection index that combined GEBVs (Torres et al., 2019; Andrade, Sousa, Oliveira, et al., 2019b). Selected parents were arranged in a partially factorial crossing scheme to maximize parental combinations and to produce sufficient within-family segregation for selection. Crosses were planned at random among the clones composing each crossing block. Whenever possible, priority was given to crosses between clones originating from different progeny in order to maximize genetic diversity within families and increase segregation potential. However, in many instances, crossing decisions were constrained by the flowering capacity of individual clones during the six-month crossing period. As a result, some progenies were derived from crosses between half-sib individuals.

Seedlings originating from the planned crosses were established in the crossing blocks and genotyped at the nursery stage (early genotyping) using DArTSeq technology. The same genotyping procedures, genomic prediction models, GEBV estimation, and parental selection criteria for the subsequent recurrent selection cycle used in the C0 phase were also adopted at this stage.

In parallel with parental selection for population improvement, the best cassava clones were evaluated across a network of multi-environment trials using the same experimental designs described for C0. Phenotypic data were processed using mixed-model approaches to estimate genotype effects, generating BLUEs and BLUPs that were incorporated into the training set for the next cycle of genomic model calibration.

#### Formation of Cycles 2 and 3 (C2 and C3)

Recurrent selection cycles C2 and C3 followed the same operational framework established for C1, with two key enhancements: (i) continuous expansion of the training population through the inclusion of newly evaluated C1 phenotypic data, and (ii) recalibration of genomic prediction models to account for shifts in allele frequencies and accumulated selection histoAt each inter-cycle transition, genomic models were retrained using the most recent phenotypic and genotypic datasets. Predictive performance was reassessed using cross-validation strategies that explicitly preserved temporal structure and family relationships, reflecting realistic breeding decision contexts.

Given the documented importance of dominance and directional dominance effects for cassava yield components, additive–dominant models were used to support population improvement and also clone selection. Model comparison, validation procedures, and selection criteria closely followed the methodological framework validated by Andrade, Sousa, Wolfe, et al. (2022).

### Phenotypic traits

A comprehensive set of traits relevant to the continuous genetic improvement of yield and root quality was evaluated throughout the recurrent selection cycles. The assessed traits included:

1. Dry Matter Content (DMC): estimated using the gravimetric method described by Kawano, Fukuda, and Cenpukdee (1987). Approximately 5 kg of clean roots were weighed in air, then in water, after excess soil was removed and the root ends were trimmed. Dry matter content was then calculated as: 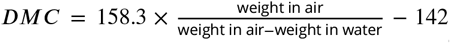, where weight in air and weight in water correspond to the root weights measured in air and water, respectively. DMC is expressed in %;
2. Starch content (Starch grav), obtained by specific weight according to Kawano, Fukuda, and Cenpukdee (1987) and expressed as a percentage;
3. Commercial fresh root yield (C.FRY): measured in *t*⋅*ha*^−1^ and defined as the yield of fresh roots free from pest and disease symptoms and with genotype-specific standard size and shape suitable for commercial use;
4. Non-commercial fresh root yield (NC.FRY): measured in *t* ⋅*ha*^−1^ and corresponding to the yield of fresh roots that do not meet commercial quality standards, representing the complement of C.FRY;
5. Total fresh root yield (FRY): calculated as the sum of commercial (C.FRY) and non-commercial (NC.FRY) fresh root yields and expressed in *t* ⋅ *ha*^−1^;
6. Dry root yield (DRY): measured in *t* ⋅ *ha*^−1^ and obtained by multiplying FRY by DMC;
7. Shoot yield (SHY): determined as total aboveground biomass, including leaves, petioles, and stems, and expressed in *t* ⋅ *ha*^−1^;
8. Harvest index (HI): calculated as 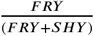 and expressed as a percentage;
9. Plant architecture (Pl.Arc): assessed on a 1–5 visual scale, where 5 indicates excellent architecture (non-branched plants or erect stems), 4 good architecture (branching above 1.60 m or low branching with at least 1.60 m of erect stem), 3 moderate architecture (branching above 1.20 m or low branching with at least 1.20 m of erect stem), 2 poor architecture (branching above 0.80 m or low branching with at least 0.80 m of erect stem), and 1 very poor architecture (highly branched clones with less than 0.80 m of erect stem);
10. Early plant vigor (Vigor): evaluated at 1.5 months after planting using a 1–5 scale, where 1 indicates low vigor, 3 intermediate vigor, and 5 high vigor;
11. Average number of roots per plant (NRP);
12. Plant height (PH), measured manually from the ground to the plant’s meristem, expressed in meters (m);
13. Number of stems per plant (NSP);

### Genotypic data

DNA extraction and quality control were performed at Intertek (Australia), followed by the construction of genomic libraries using the DArTseq complexity-reduction method (Kilian et al., 2012) at Diversity Arrays Technology (Canberra, Australia). Sequencing was conducted on the Illumina HiSeq 2500 platform (Illumina, USA). Raw data processing involved barcode removal and trimming using STACKS software (Catchen et al., 2013). Reads were aligned to the Cassava Reference Genome v 6.1 using BWA (Li, 2010). Variant calling was performed with GATK (Auwera and O’Connor, 2020), excluding indels and multiallelic sites. Quality filtering used TASSEL GBS 5.0 pipeline v 5.2.90 (Glaubitz et al., 2014) with filters: minimum call rate 80%, minimum allele frequency (MAF) 1%, and imputation by LD KNNi method (Money et al., 2015). The final dataset contained 25,923 informative SNPs.

### Phenotypic data analysis

For this study, a total of 210 field trials conducted between 2011 and 2024 were analyzed. The trials were distributed across multiple cities and implemented using either a randomized complete block design or an augmented block design. The evaluated accessions represented multiple stages of the breeding pipeline, including improved varieties derived from controlled crosses, materials originating from recurrent selection cycles (advanced populations), and a limited number of varieties sourced from research institutions or identified by farmers, which served as reference germplasm. Across all analyses, 23 accessions were used as checks, rotated among trials without maintaining a fixed proportion across experiments.

#### Single-trial and multi-trial analyses

The experimental dataset exhibited substantial imbalance in both experimental design and population composition across and within trials. In response to this structure, a two-stage phenotypic analysis was adopted. In the first stage, phenotypic data were analyzed separately for each trial to account for experimental design effects and to remove non-genetic sources of variation. BLUEs were obtained for each accession within each environment (combination between location and year) using the following linear mixed model:

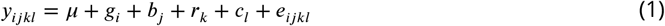

where *y*_*ijkl*_ is the plot observation for genotype *i* at the block *j*, row *k*, and column *l*; *μ* is the overall mean; *g*_*i*_ is the fixed effect of the *i*_*th*_ genotype; *b*_*j*_ is the random effect of the *jth* block; *r*_*k*_ and *c*_*l*_ are the random effects of the *k*_*th*_ row and *l*_*th*_ column, respectively. Row and column effects were modeled using a penalized tensor product of marginal B-splines, following Rodríguez-Álvarez et al. (2018). Residuals (*e*_*ijkl*_) were assumed to be independent and identically distributed.

In the second stage, the trial-specific BLUEs were used as input data in mixed models to estimate genotype BLUEs across environments for each evaluated trait. The following models were fitted:

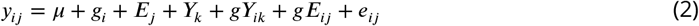

where *y*_*ij*_ is the phenotypic value of the trait for the *i*_*th*_ genotype evaluated in the *j*_*th*_ environment; *μ* is the overall mean; *g*_*i*_ is the fixed effect of the *i*_*th*_ genotype; *Y*_*k*_ is the fixed effect of the *k*_*th*_ year; *E*_*j*_ is the random effect of the *j*_*th*_ environment; *gY*_*ik*_ is the random effect of the interaction between *i*_*th*_ genotype and the *k*_*th*_ year; *gE*_*ij*_ represents the random effect of the interaction between the *i*_*th*_ genotype and the *j*_*th*_ environment and *e*_*ij*_ is the residual error term.

Residual variances were assumed to be heterogeneous across environments, following a multivariate normal distribution *e*_*ijk*_ ∼ *N*(0, *W* ), where W is a diagonal matrix containing the inverse of the standard errors of the BLUEs from the first stage 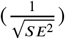. These standard errors were used to appropriately scale the residual variance in the second-stage analysis, as described by Frensham, Cullis, and Verbyla (1997) and Smith, Cullis, and Gilmour (2001).

In the first stage, outliers were additionally identified and removed using the Bonferroni-adjusted outlier test, implemented via the `outlierTest` function in the `car R` package (Fox and Weisberg, 2018). In the second stage, the final mixed-model structure was determined based on the statistical significance of random effects. Random terms were evaluated using Likelihood Ratio Tests (LRTs) by comparing the full model against reduced models obtained by sequentially removing random effects, with inference based on the chi-square distribution.

Following the estimation of genotype BLUEs, genotypes were subsequently refitted as random effects to obtain the genetic variance component. This estimate was then used to calculate the phenotypic heritability (*H*^2^) for each trait, following Cullis, Smith, and Coombes (2006):

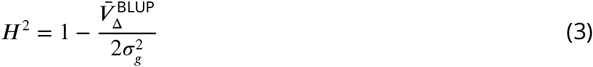

Phenotypic accuracy (*ACC*) for each trait was computed as:

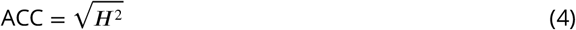

where 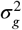 denotes the genotypic variance, 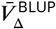 is the mean variance of the difference between two genotype BLUPs. All phenotypic analyses were performed using ASReml-R version 4.2 (Butler et al., 2009).

### Genomic best linear unbiased prediction (GBLUP)

The consistency of genomic selection performance across recurrent selection cycles was evaluated using historical phenotypic and genotypic data from the breeding program, analyzed separately for each trait. For each cycle–trait combination, independent phenotypic datasets and corresponding genomic relationship matrices were constructed. Historical information was incorporated incrementally, such that only phenotypic records available up to a given selection cycle were used in the analysis.

For cycle C0, the dataset comprised 25 multi-environment trials conducted between 2011 and 2016. Cycle C1 included phenotypic data from 31 trials evaluated between 2011 and 2018, which included the 25 trials of C0 and six additional trials derived from population updates. Cycle C2 consisted of 50 trials conducted between 2011 and 2020, including all trials of C0 and C1, supplemented by 19 newly established update trials. Finally, cycle C3 incorporated 160 additional trials beyond those included in earlier cycles. This dataset included trials evaluating clones derived from both previous genomic selection cycles and the program’s base population, the latter consisting of newly introduced germplasm originating from other breeding programs, evaluation nurseries, or other external sources incorporated into the breeding pipeline.

For each genomic selection cycle, predictive performance was assessed via cross-validation using 100 independent replicates. In each replicate, clones were randomly partitioned into a training set comprising 80% of individuals with both genotypic and phenotypic information, and a validation set consisting of the remaining 20% of individuals, for which only genotypic data were available.

To evaluate the internal predictive consistency of each genomic selection cycle within its corresponding historical dataset, the GBLUP model was fitted independently for each trait and cycle according to the following linear mixed model:

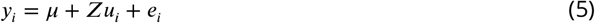

where *y*_*i*_ is the vector of BLUEs for trait *i* ; *μ* is the overall mean; *Z* is the incidence matrix relating observations to genomic effects; 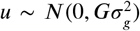 is the vector of additive genomic breeding values, with *G* denoting the genomic relationship matrix constructed following VanRaden (2008) and 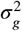 the additive genetic variance and 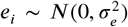 is the residual error, where 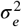 represents the residual variance.

The models were implemented using the BGLR package (Pérez-Rodríguez and Campos, 2022). A total of 22,000 MCMC iterations were performed, including 2,000 burn-in iterations.

Model performance was evaluated using the following metrics:

Predictive ability (PA), defined as the Pearson correlation between predicted GEBVs in the validation set and the observed phenotypic BLUEs, and Root Mean Square Error (RMSE), used to quantify prediction precision:

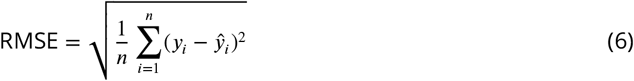

where *n* represents the number of observations, *y*_*i*_ is the observed value, and *ý*_*i*_ is the predicted value.

### Realized genetic gain across recurrent genomic selection cycles

GEBVs obtained from the GBLUP model were used to quantify RGG across recurrent selection cycles. For this purpose, GEBVs derived from the final evaluation cycle were used, as this dataset integrates cumulative phenotypic and genotypic information from three successive selection cycles, thereby maximizing the precision and comparability of genomic predictions across cycles. For each trait, clone-specific GEBVs were analyzed using linear regression models in which genomic values were regressed on the selection cycle. Each clone was assigned to the selection cycle corresponding to its first field evaluation, reflecting its initial introduction into the breeding pipeline and ensuring temporal consistency in estimating genetic trends.

Realized genetic gain was estimated as the slope of the regression of GEBVs on selection cycles, representing the average change in genetic merit per selection cycle. To express genetic gain on a relative scale and enable comparisons across traits, the following percentage-based metric was calculated:

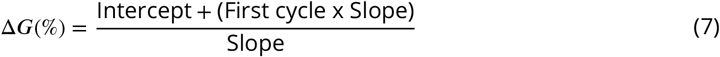

This formulation allows quantification of relative trait improvement across cycles, enabling direct comparisons of genomic selection efficiency across traits, cycles, and temporal scales (per cycle or per year) (Covarrubias-Pazaran, 2020; Seck et al., 2023; Rutkoski, 2019a).

### Assessment of genetic diversity within and across selection cycles

Genetic diversity parameters within and between selection cycles were evaluated using a set of complementary metrics. Initially, the genotypic data were partitioned into subsets containing only clones from each selection cycle, thereby allowing characterization of within-cycle genetic diversity. Subsequently, the complete dataset was used to investigate overall patterns of genetic diversity across the entire breeding process represented by the population under study.

To assess genetic diversity within each selection cycle, observed heterozygosity (*H*_*o*_) and the individual inbreeding coefficient (*F*_*i*_) were estimated:

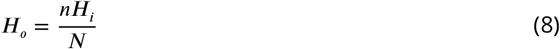

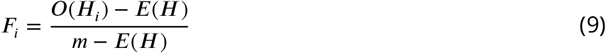

where *nH*_*i*_ is the number of heterozygous genotypes (*A*_1_*A*_2_ or *A*_2_*A*_1_) observed in individual *i, N* is the total number of markers evaluated, *m* is the number of genotyped markers, *O*(*H*_*i*_) is the observed heterozygosity of individual *i*, and *E*(*H*) = ∑_*j*_ [1 − 2*p*_*j*_(1 − *p*_*j*_)] represents the expected homozygosity across all SNP markers.

Genetic diversity among selection cycles was quantified using Nei’s genetic distance (*GD*) (Nei, 1972; Nei, 1978) and the pairwise fixation index (*F*_*ST*_ ) (Wright, 1949):

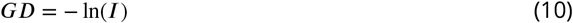

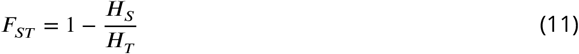

where *I* denotes the genetic identity between pairs of populations, *H*_*S*_ is the average expected heterozygosity within subpopulations, and *H*_*T*_ is the expected heterozygosity in the total population.

The statistics *H*_*o*_, *F*_*i*_, and *F*_*ST*_ were estimated using the `snpReady` package (Granato et al., 2018), whereas Nei’s genetic distance (*GD*) was calculated using the `adegenet` package (Jombart and Ahmed, 2011).

## Results

### Phenotypic variation, genetic parameters and trait reliability across cycles

Across the 210 multi-environment trials conducted between 2011 and 2024, substantial phenotypic variation was observed for all evaluated agronomic and quality traits. Traits associated with biomass accumulation and root production exhibited the greatest phenotypic dispersion. The highest phenotypic means were observed for harvest index, fresh root yield, commercial fresh root yield, shoot yield, dry matter content, and starch content. In contrast, physiological and architectural traits, including early plant vigor, number of roots per plant, and number of stems per plant, displayed lower phenotypic means and reduced variation (Table 1).

**Table 1.** Phenotypic means (*μ*), standard error (SE), phenotypic accuracy (ACC), and heritability (*H*^2^) for agronomic and physiological traits assessed across 13 years of multi-environment cassava breeding trials targeting enhanced root industrial quality.

| Trait | $\mu$ | SE | ACC | $H^2$ |
| --- | --- | --- | --- | --- |
| Dry matter content (%) | 35.22 | 2.64 | 0.78 | 0.61 |
| Dry root yield ($t \cdot ha^{-1}$) | 8.93 | 2.99 | 0.69 | 0.48 |
| Commercial fresh root yield ($t \cdot ha^{-1}$) | 19.12 | 10.82 | 0.64 | 0.41 |
| Non-commercial fresh root yield ($t \cdot ha^{-1}$) | 5.47 | 3.84 | 0.50 | 0.25 |
| Total fresh root yield ($t \cdot ha^{-1}$) | 24.65 | 11.92 | 0.68 | 0.46 |
| Harvest index (%) | 41.57 | 14.32 | 0.74 | 0.55 |
| Number of roots per plant | 4.39 | 2.40 | 0.51 | 0.26 |
| Number of stems per plant | 2.01 | 0.64 | 0.59 | 0.34 |
| Plant height (m) | 2.22 | 0.49 | 0.77 | 0.60 |
| Plant architecture (1 to 5 visual scale) | 2.57 | 1.18 | 0.73 | 0.54 |
| Shoot yield ($t \cdot ha^{-1}$) | 30.69 | 12.45 | 0.71 | 0.51 |
| Starch content (m) | 20.50 | 2.88 | 0.78 | 0.61 |
| Early plant vigor (1 to 5 visual scale) | 2.16 | 0.90 | 0.43 | 0.18 |

Across the 210 multi-environment trials conducted between 2011 and 2024, substantial phenotypic variation was observed for all evaluated agronomic and quality traits. Traits associated with biomass accumulation and root production exhibited the greatest phenotypic dispersion. The highest phenotypic means were observed for HI, FRY, FRY.C, SHY, DMC, and starch content. In contrast, physiological and architectural traits, including early plant vigor, NRP, and NSP, displayed lower phenotypic means and reduced variation (Table 1).

Genetic parameter estimates revealed clear differences among trait categories. Moderate to high broad-sense heritability was estimated for structural, compositional, and biomass-partitioning traits, including plant height, plant architecture, HI, DMC, starch content, and SHY, with values ranging from approximately 0.45 to 0.61 (Table 1). These traits also exhibited moderate to high ACC, indicating reliable genetic evaluation across environments.

In contrast, traits related to early plant vigor and root number presented low heritability (< 0.20) and reduced ACC, suggesting lower precision of phenotypic estimates under the current experimental design. Among all evaluated traits, early plant vigor exhibited the lowest ACC. Overall, traits associated with root yield, dry matter accumulation, and biomass partitioning consistently displayed the most favorable genetic properties, supporting their use in subsequent genomic analyses.

### Predictive ability dynamics and prediction error across recurrent genomic-selection cycles

PA and RMSE were evaluated across four recurrent genomic selection populations spanning 2011 to 2024, under multiple training–target population scenarios. These scenarios were designed to mimic operational breeding decisions, ranging from predictions between temporally adjacent cycles to predictions targeting more advanced cycles using expanded historical training sets.

The PA of the GBLUP models varied among traits and across recurrent genomic selection cycles (Figure 1). Across all cycles, PA values ranged from low (≥ 0.30) to moderate-to-high (*≈* 0.59), depending on trait genetic architecture and data availability. Plant height emerged as the most consistently predictable trait across all cycles, with PA values remaining between 0.49 and 0.55 in C0, C1, C2, and C3. Similarly, starch gravimetric content and DMC showed stable and moderate predictive ability across cycles, particularly in C1 and C2, where PA frequently exceeded 0.40.

**Figure 1.**
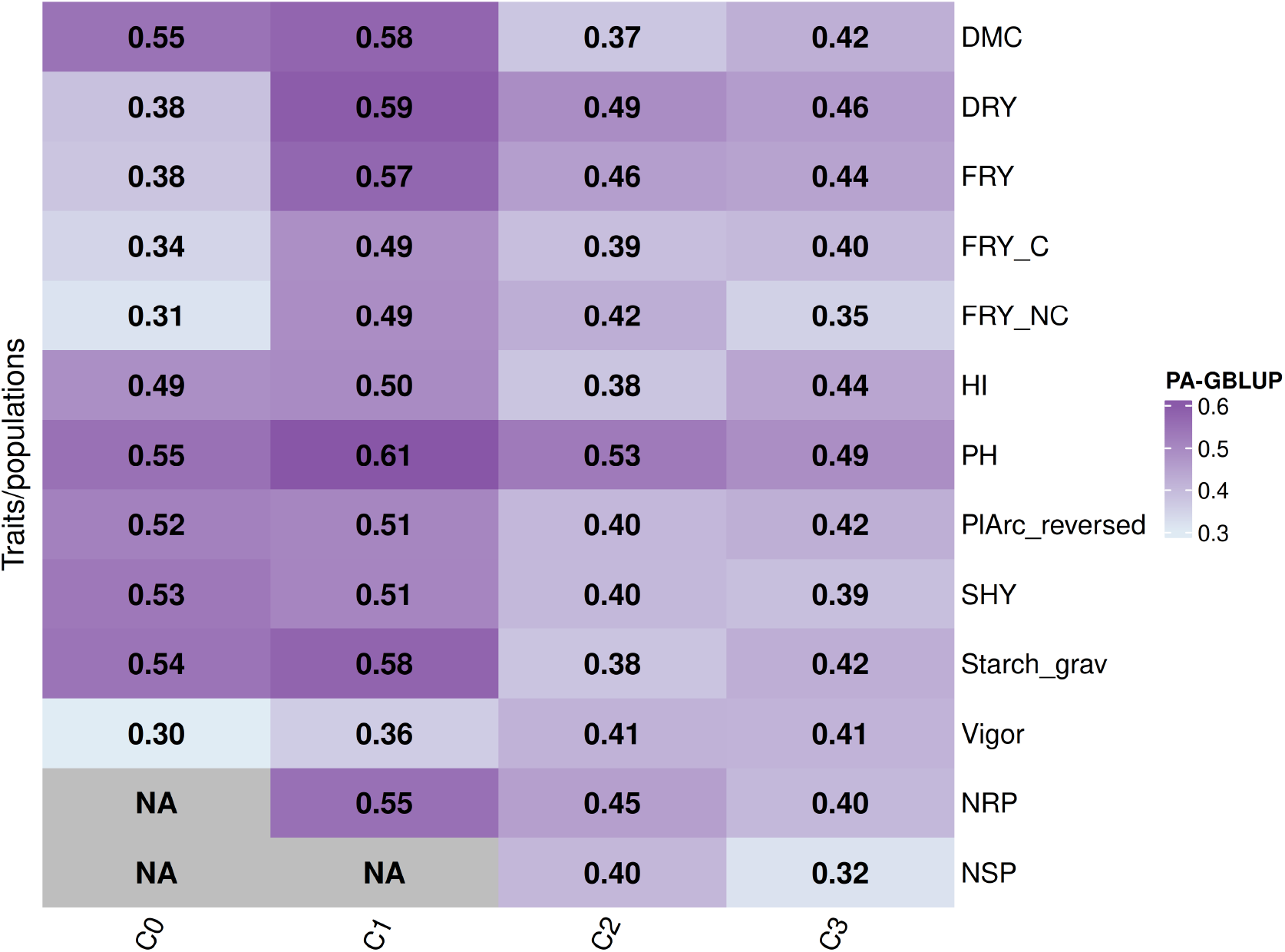
Heatmap of Predictive Ability for agronomic and quality traits across four training–target population scenarios in terms of GS populations (C0, C1, C2, and C3) spanning the 2011–2024 evaluation period. Darker shades denote higher prediction accuracy, whereas NA values indicate unavailable or non-estimable scenarios. *Abreviations*: DMC:Dry matter content; Starch grav: Starch content; C.FRY: Commercial fresh root yield; NC.FRY:Non-commercial fresh root yield;FRY: Total fresh root yield; DRY: Dry root yield; SHY: Shoot yield; HI: Harvest index; PlArc: Plant architecture; Vigor: Early plant vigor; NRP:Average number of roots per plant; NSP: Plant height PH;Number of stems per plant.

**Figure 2.**
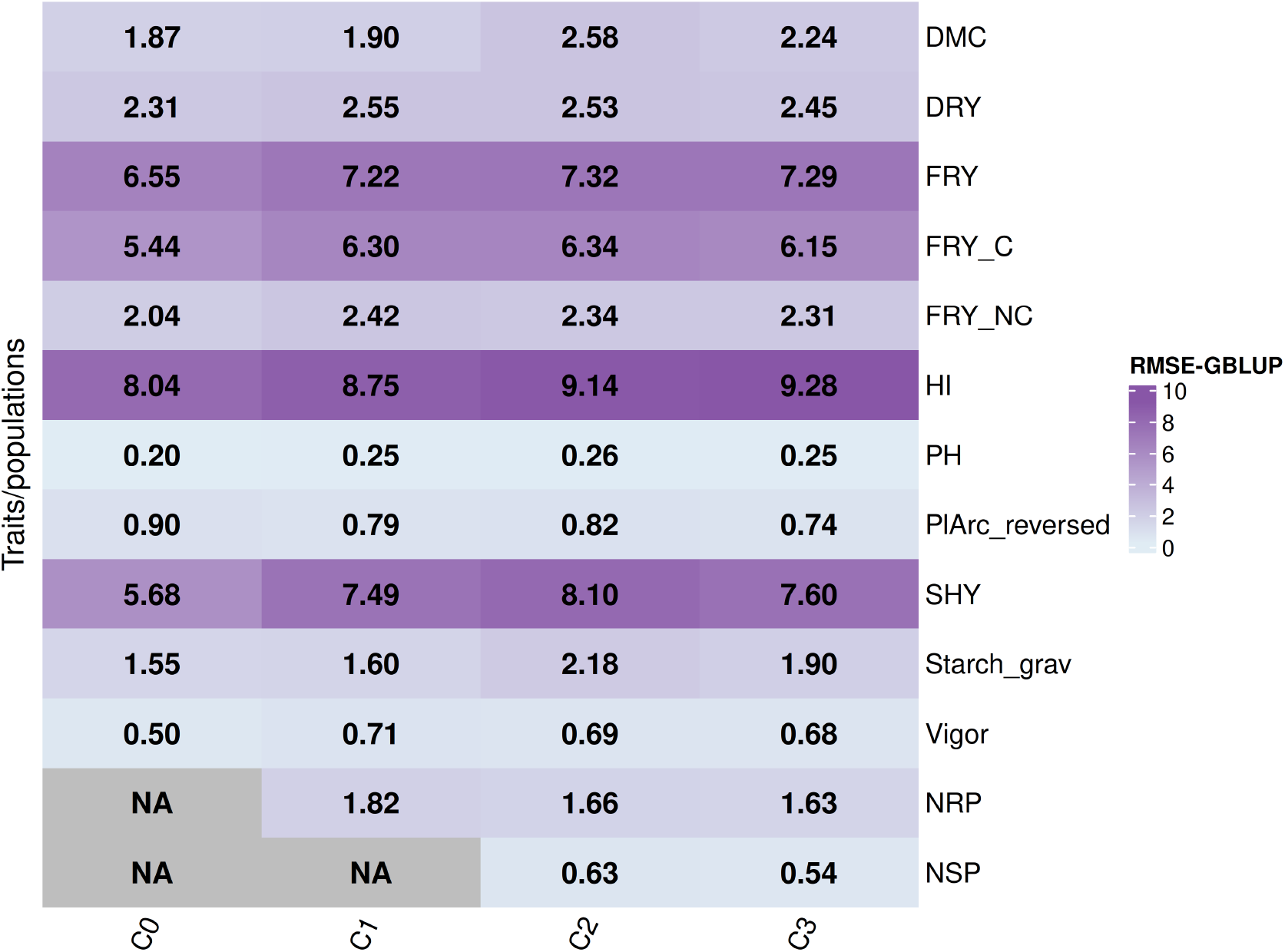
Heatmap of RMSE for agronomic, quality traits across four training–target population scenarios in terms of GS populations (C0, C1, C2, and C3) spanning the 2011–2024 period. Darker shades denote higher prediction error, whereas NA values indicate unavailable estimates. *Abreviations*: DMC:Dry matter content; Starch grav: Starch content; C.FRY: Commercial fresh root yield; NC.FRY:Non-commercial fresh root yield;FRY: Total fresh root yield; DRY: Dry root yield SHY: Shoot yield, HI: Harvest index; PlArc: Plant architecture: Vigor: Early plant vigor; NRP:Average number of roots per plant; NSP: Plant height PH;Number of stems per plant.

Yield-related traits, including DRY, FRY, and SHY, exhibited moderate PA values, typically ranging between 0.38 and 0.59. Predictive ability for these traits was highest when training and validation populations were temporally adjacent (e.g., C0 to C1, C1 to C2) and declined modestly when predictions targeted more advanced cycles. However, when cumulative historical data were incorporated into the training populations (C2 and C3), PA values stabilized or improved compared with single-cycle training scenarios.

Some Physiological or architectural traits, including early plant vigor, plant architecture, and number of roots per plant, consistently displayed moderate PA (between 0.30 and 0.45 across several scenarios). Increasing the training population size across cycles yielded only marginal improvements in these traits, and predictive performance remained comparatively limited even in later cycles. Overall, predictive ability was highest in C1, intermediate in C0, and showed a slight decline in C2 and C3 for several traits. Nevertheless, for most agronomically relevant traits, PA remained within operationally useful ranges throughout all four selection cycles.

RMSE estimates complemented PA results by quantifying prediction precision across traits and cycles (Figure 1). Yield-related traits such as FRY, DRY, SHY, and HI exhibited the highest RMSE values, whereas compositional traits, including DMC and starch content, showed consistently low RMSE across all cycles. For most traits, RMSE declined moderately as training populations expanded from C0 to C3, indicating improved model calibration as historical phenotypic information accumulated. This trend was particularly evident for NRP, NSP, Early plant vigor, and Plant architecture, where RMSE decreased in C2 and C3 compared with earlier cycles. Physiological traits such as early vigor and plant architecture exhibited relatively stable RMSE values across cycles and showed limited sensitivity to training population expansion. Taken together, RMSE and PA patterns demonstrate that while absolute prediction error remained trait-dependent, genomic prediction models maintained consistent precision across recurrent selection cycles.

### Realized genetic gain across recurrent genomic-selection cycles

Regression analyses of clone-specific GEBVs on selection cycle revealed sustained genetic change across the 13-year period of genomic selection (Table 2). For most traits, regression slopes differed significantly from zero, indicating measurable realized genetic gain across cycles. Among yield-related traits, the largest positive gains per cycle were observed for dry root yield (DRY; slope = 0.44, *p* − *value* < 0.01), shoot yield (SHY; slope = 1.21, *p* − *value* < 0.01), early plant vigor (slope = 0.39, *p* − *value* < 0.01), number of stems per plant (NSP; slope = 0.05, *p* − *value* < 0.01), and plant height (PH; slope = 0.03, *p* − *value* < 0.01). When expressed on a relative scale, cumulative genetic gains for most yield-related traits ranged from approximately 1.5% to 5% across four selection cycles. Early plant vigor showed the largest relative increase, with 18,47 % over the evaluated period.

**Table 2.** Realized genetic gain in percentage (RGG %) based on GEBVs for agronomic and quality traits across four recurrent genomic selection cycles (C0 to C3) conducted between 2011 and 2024. The regression slope represents the average genetic change per cycle.

| Trait | Slope | $p$-value | Intercept | RGG (%) |
| --- | --- | --- | --- | --- |
| Dry matter content (%) | -0.03 | 0.37 | 35.29 | -0.08 |
| Dry root yield ($t \cdot ha^{-1}$) | 0.47 | < 0.01 | 8.82 | 5.41 |
| Commercial fresh root yield ($t \cdot ha^{-1}$) | -0.12 | 0.04 | 19.46 | -0.60 |
| Non-commercial fresh root yield ($t \cdot ha^{-1}$) | 0.15 | < 0.01 | 5.34 | 2.86 |
| Total fresh root yield ($t \cdot ha^{-1}$) | 0.13 | 0.15 | 25.14 | 0.44 |
| Harvest index (%) | -1.72 | < 0.01 | 43.41 | -3.97 |
| Number of roots per plant | -0.17 | < 0.01 | 5.48 | -3.59 |
| Number of stems per plant | 0.05 | < 0.01 | 2.24 | 2.51 |
| Plant height (m) | 0.03 | < 0.01 | 2.18 | 1.51 |
| Plant architecture (1 to 5 visual scale) | -0.07 | < 0.01 | 2.66 | -2.62 |
| Shoot yield ($t \cdot ha^{-1}$) | 1.21 | < 0.01 | 30.12 | 4.03 |
| Starch content (%) | -0.02 | 0.34 | 20.57 | -0.12 |
| Early plant vigor (1 to 5 visual scale) | 0.39 | < 0.01 | 2.10 | 18.47 |

In contrast, negative trends were detected for traits associated with quality and plant architecture, including dry matter content (DMC),fresh root yield (FRY.C), harvest index (HI), number of roots per plant (NRP), starch content (Starc Grav), and plant architecture (Pl.Arc). For DMC and Starch Grav, although the regression slopes were negative, these effects were not statistically significant. The modest declines observed for DMC and starch content are biologically consistent with the well-established negative genetic correlation between starch concentration and fresh root yield in cassava. Importantly, the magnitude of these reductions was limited (−0.07% to −0.11% per cycle), indicating that yield gains were achieved without substantial deterioration of root quality. For FRY, a positive but non-significant slope was observed, resulting in only minor relative gains; this pattern may have indirectly affected correlated traits such as FRY.C, HI, and NRP. Nevertheless, the observed declines in these traits remained modest (< 4%), suggesting a limited impact on overall productive performance.

For plant architecture, the negative trend (PlArc; slope = 0.05) reflects intentional selection toward lower scores, corresponding to more erect and less branched plant types that are better suited for mechanization. Overall, these results indicate that genomic selection promoted consistent genetic progress for yield and vigor traits, while revealing predictable trade-offs with correlated quality and architectural traits that can be effectively managed over time through balanced selection indices and refined breeding objectives.

### Population genetics, diversity erosion, and structural shifts across recurrent genomic selection cycles

Genetic diversity and population structure analyses revealed gradual changes in the population’s genetic composition across genomic selection cycles (Table 3; Figure 3). Both the proportion of genetic variation attributable to differences among populations *F*_*ST*_ and the genetic divergence based on allele frequencies (Nei’s genetic distance) indicated consistent patterns of differentiation throughout the selection process.

**Table 3.**
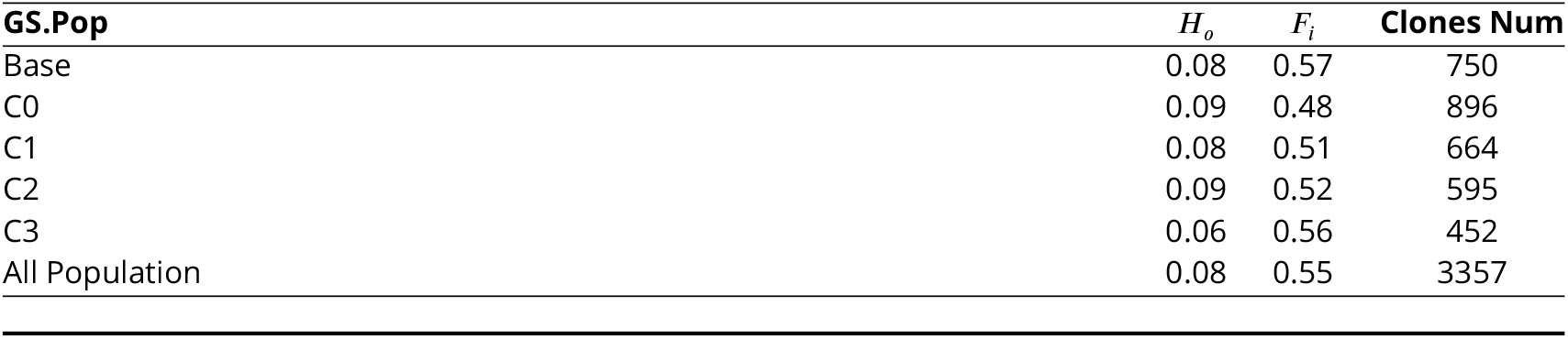
Genetic diversity statistics estimated for each recurrent genomic selection cycle (C0 to C3) for the combined population, including observed heterozygosity (*H*_*o*_), inbreeding coefficient (*F*_*i*_), and the number of evaluated clones per cycle (*ClonesN um*).

| GS.Pop | $H_o$ | $F_i$ | Clones Num |
| --- | --- | --- | --- |
| Base | 0.08 | 0.57 | 750 |
| C0 | 0.09 | 0.48 | 896 |
| C1 | 0.08 | 0.51 | 664 |
| C2 | 0.09 | 0.52 | 595 |
| C3 | 0.06 | 0.56 | 452 |
| All Population | 0.08 | 0.55 | 3357 |

**Figure 3.**
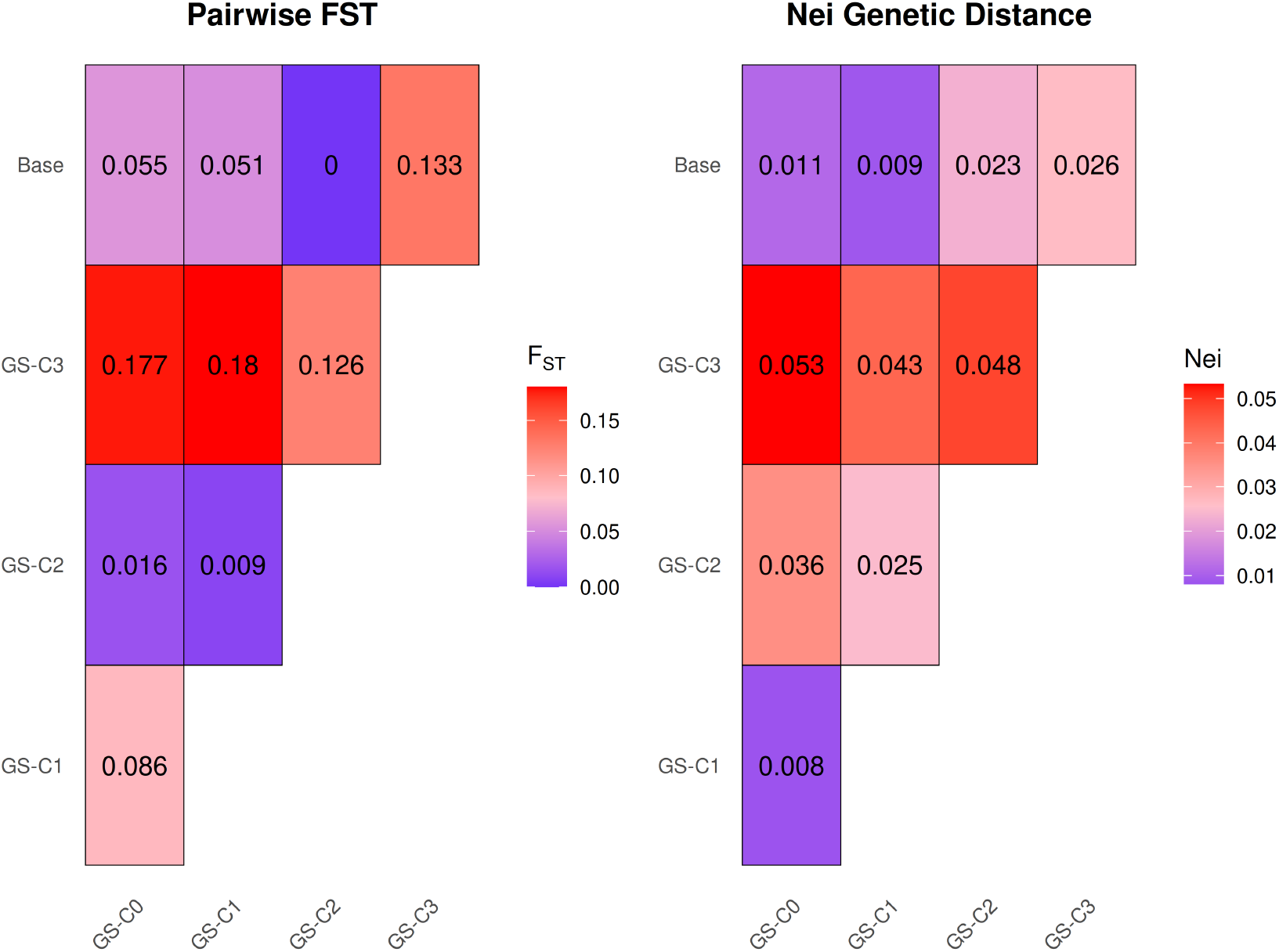
Pairwise genetic differentiation (*F*_*ST*_ ; left panel) and Nei’s standard genetic distance (*GD*; right panel) among recurrent genomic selection cycles (C0 to C3), estimated using genome-wide SNP markers.

Overall, low levels of genetic divergence were observed between the Base population and the C0, C1, and C2 cycles, with no evidence of substantial population subdivision among these groups. Comparisons between C1 and C2 showed the lowest levels of genetic differentiation (*F*_*ST*_ ≤ 0.05; Nei’s genetic distance < 0.05), indicating a high degree of genetic similarity between these cycles.

In contrast, the C3 population exhibited the highest levels of differentiation relative to the other populations (*F*_*ST*_ ≥ 0.10; Nei’s genetic distance ≥ 0.04), suggesting a greater degree of genetic divergence accumulated over successive selection cycles. Relative to the Base population, C3 showed an approximately 25% reduction in observed heterozygosity, accompanied by changes in inbreeding patterns (Table 3).

The observed heterozygosity (*H*_*o*_) and inbreeding coefficient (*F*_*i*_) estimates further supported the patterns identified by the *F*_*ST*_ and Nei’s genetic distance analyses. The Base, C0, C1, and C2 populations displayed relatively similar levels of observed heterozygosity and inbreeding (*H*_*o*_ = 0.08–0.09; *F*_*i*_ = 0.48–0.56), reinforcing the genetic similarity among these groups. In contrast, C3 exhibited the lowest Ho values, particularly when compared with C0 and C2. Inbreeding coefficients were relatively high across all populations, ranging from 0.48 to 0.57, with the highest values observed in the Base population (*F*_*i*_=0.57) and C3 (*F*_*i*_=0.55). The combination of reduced observed heterozygosity and elevated inbreeding suggests that the cumulative effects of selection became more evident in C3, reflecting a reduction in the available genetic diversity within this population.

Nevertheless, some factors should be considered when interpreting these results. First, C3 included fewer clones than the previous cycles, which may have increased sampling variance in the estimates of heterozygosity and inbreeding. Second, operational constraints in the crossing blocks reduced seed production in the originally planned recombination year, leading to the division of the C3 population into two batches, with the second established one year later. This temporal separation reduced the number of individuals available during the originally planned evaluation stage and may have influenced estimates of effective population size and the genetic diversity parameters observed for this cycle.

## Discussion

Genetic gain in plant breeding is fundamentally determined by the joint optimization of selection accuracy, selection intensity, genetic variance, and breeding cycle length. Genomic selection directly targets three of these components by increasing prediction accuracy, enabling the capture of genome-wide additive variance, and substantially shortening breeding cycles, even under comparable selection intensities to phenotypic selection (Crossa et al., 2017; Das, Vinayan, Seetharam, et al., 2021; Wartha and Lorenz, 2021). Consequently, GS has shifted its focus from expected to realized genetic gain as the most informative metric for evaluating program efficiency, sustainability, and long-term genetic progress, with RGG integrating biological response, operational constraints, and population-level dynamics (Rutkoski, 2019b; Seck et al., 2023).

The present study represents a comprehensive effort to quantify realized genetic gain under recurrent genomic selection within the Brazilian cassava breeding program. By integrating historical phenotypic data, genomic prediction, and population genetic analyses across four recurrent selection cycles spanning more than a decade, this work provides a robust framework to evaluate not only short-term predictive performance but also long-term responses and changes in population structure. Such integrative assessments remain scarce in clonally propagated crops, where the balance between rapid genetic gain and maintenance of diversity is particularly critical (Rutkoski, 2019b; Zheng et al., 2022).

### Heritabilities, genetic architecture, and implications for phenotyping

The two-stage phenotypic modeling framework employed in this study demonstrated good internal coherence in partitioning genetic and environmental variance across productive, physiological, and architectural traits. Traits exhibiting moderate to high heritability consistently showed higher genomic prediction accuracy, reinforcing the fundamental principle that additive genetic variance is the primary driver of predictive performance under GS (Daetwyler et al., 2010; Crossa et al., 2017).

Most production-related and physiological traits displayed heritability estimates in the range of 0.25—0.60, consistent with expectations for complex quantitative traits in cassava. These values closely align with previous estimates reported for Brazilian germplasm, including dry matter content, fresh root yield, shoot yield, plant height, and harvest index (Andrade, Sousa, Oliveira, et al., 2019b; Sampaio Filho et al., 2023b; Costa et al., 2024). In contrast, traits such as early vigor, number of roots per plant, and Non-commercial fresh root yield exhibited low heritability (≤ 0.25), reflecting their strong sensitivity to micro-environmental variation, developmental stage, and management practices. This effect was particularly evident for physiological traits, which are known to exhibit pronounced environmental sensitivity in cassava (Manze et al., 2021). These results highlight the need for complementary evaluation strategies, repeated measures over time, and standardized high-throughput phenotyping platforms. Empirical evidence indicates that such approaches can substantially increase heritability estimates and enhance the detection of genetic signals for complex traits (Nascimento, Cortes, et al., 2025).

### Dynamics of predictive performance across selection cycles

The predictive accuracies obtained in this study ranged from low to moderate, reflecting intrinsic differences in genetic architecture, heritability, and signal-to-noise ratio among traits and breeding populations. Consistent with previous reports, variation in predictive ability across productive and physiological traits was primarily driven by differences in trait heritability and the extent to which additive genetic variance could be captured by genome-wide markers (Berro et al., 2019). In this context, productive traits such as fresh root yield, as well as physiological and architectural traits including plant height, plant architecture, shoot yield, and number of stems per plant, consistently exhibited higher predictive accuracy and greater reliability across selection cycles.

Beyond predictive accuracy, a key pattern emerging from the results was stable or progressively reduced RMSE values across the more advanced selection cycles, with no evidence of abrupt increases. This trend indicates a gradual improvement in prediction precision over successive cycles. Notably, the reduction in RMSE occurred even when predictive accuracy did not increase proportionally, suggesting that the genomic prediction models became better calibrated over time. These findings indicate that, despite increasing genetic complexity and continued recombination throughout the recurrent breeding process, prediction error remained controlled rather than accumulating. Overall, this pattern reinforces the efficiency and robustness of genomic selection under recurrent breeding schemes.

The influence of selection cycle advancement and training population composition, particularly the contribution of the base population (C0), was evident in the temporal trajectory of predictive performance. An initial increase in predictive accuracy from C0 to C1 was followed by a slight decline in later cycles (C2 and C3), a pattern indicative of increasing genetic distance between training and validation populations. Such decay in predictive ability is a well-documented consequence of directional selection and recombination, which progressively alter allele frequencies and linkage disequilibrium patterns (Das, Vinayan, Patel, et al., 2020; Das, Vinayan, Seetharam, et al., 2021). The higher predictive accuracy and realized genetic gain observed in early cycles are likely associated with the greater genetic diversity and heterozygosity present in C0–C1, whereas the gradual decline in later cycles reflects the cumulative reduction in genetic variability resulting from recurrent selection.

An additional factor contributing to reduced predictive accuracy in advanced cycles was the incorporation of accessions from outside the breeding program, beginning with cycle C2. While this strategy is beneficial for broadening the genetic base and introducing novel variation, it may also increase genetic contrast between training and selection populations, thereby reducing model transferability and prediction accuracy if not explicitly accounted for in model-updating strategies. Additionally, populations formed by combining multiple selection cycles consistently exhibited greater predictive stability, with only minor fluctuations in RMSE across traits. Our finding reinforces the principle that larger and more genetically diverse training populations enhance the robustness and generalizability of genomic prediction models, even if they do not always maximize predictive accuracy. Such stability is particularly relevant for operational breeding programs, where consistent performance across cycles is often more valuable than transient gains in accuracy.

### Genomic selection led to significant gains over a decade of evaluation

The overall success of the breeding program is evidenced by its capacity to achieve positive realized genetic gains across multiple trait groups over the 13-year evaluation period. Genetic gains were observed not only for highly heritable productive traits but also for physiological traits, which are typically more challenging to improve due to strong environmental influence and pronounced genotype × environment interaction. These results demonstrate the effectiveness of genomic selection in delivering cumulative genetic progress under realistic breeding conditions.

Negative trends observed for certain traits should be interpreted in the context of indirect selection responses and explicit breeding objectives rather than as failures of the selection strategy. Reductions in dry matter and starch content, for instance, are biologically plausible given the strong genetic correlation between these traits and the prioritization of yield-related attributes in selection indices. Similarly, the decline in plant architecture reflects intentional selection toward ideotypes better suited for mechanized production systems. From this perspective, these responses represent coherent outcomes of a multi-trait selection framework rather than unintended consequences.

Genetic gains for dry matter and starch-related traits are inherently difficult to achieve, largely due to their strong sensitivity to environmental variation and substantial G×E interaction. Previous studies have consistently reported modest rates of improvement for these traits. For example, Manze et al. (2021) observed a gain of approximately 0.10% in dry matter content for cassava clones released between 1940 and 2019 when evaluated within a single year, while Delgado et al. (2024) reported a gain of 0.53% for clones evaluated between 2013 and 2022. An important distinguishing feature of this study is the comprehensive evaluation of germplasm within each selection cycle, rather than restricting analyses to a subset of elite or advanced clones. This approach enabled a more integrative assessment of realized genetic gain across the entire breeding pipeline, capturing both short-term responses in advanced material and broader trends in population-level genetic improvement. As such, the observed gains reflect not only selection efficiency but also the cumulative impact of genomic selection on the structure and performance of the breeding population as a whole.

### Changes in population dynamics and gene flow over selection cycles

Analyses of genetic diversity indices indicate that the breeding program successfully achieved consistent genetic gains across recurrent genomic selection cycles. However, this progress occurred alongside a moderate but measurable reduction in genetic variability, a pattern widely documented in genomic selection frameworks. Because genomic selection enables shorter cycle lengths and a greater number of selection cycles per unit of time than phenotypic selection, the cumulative effects of selection and drift tend to manifest more rapidly (Rutkoski et al., 2015; Crossa et al., 2017; Gorjanc, Gaynor, and Hickey, 2018). Accordingly, successive cycles of genomic selection are frequently associated with reductions in observed heterozygosity and effective population size, coupled with increases in inbreeding coefficients.

Beyond these general trends, the present study revealed a non-linear pattern of genetic divergence among selection cycles. Genetic differentiation did not increase gradually with cycle advancement; instead, a pronounced shift was observed at cycle C3, characterized by an abrupt increase in both *F*_*ST*_ and Nei’s genetic distance. In contrast, genetic differentiation among C0, C1, and C2 remained consistently low, indicating strong genetic continuity across early cycles. This pattern strongly suggests that the observed divergence at C3 was driven primarily by demographic factors, most notably a smaller number of evaluated clones, rather than by the mere accumulation of selection cycles. Similar conclusions were reached by Gorjanc, Gaynor, and Hickey (2018), who demonstrated that genetic differentiation under genomic selection is far more sensitive to effective population size than to the number of selection cycles alone.

Breeding practices related to parental selection and mating design provide an important mechanistic explanation for these observed dynamics. Throughout most cycles, parental selection for crossing blocks relied predominantly on genotypes derived from within the breeding program and selected for high genomic estimated breeding values, with limited introgression of external germplasm and no use of optimized mating strategies. Comparable approaches have been reported in wheat and barley breeding programs, where recurrent recombination among related individuals resulted in progressive reductions in genetic diversity across cycles (Rutkoski et al., 2015; Allier et al., 2019). While such strategies are highly effective for maximizing short-term genetic gain, they may inadvertently accelerate genetic erosion if not complemented by explicit diversity management measures.

Operational constraints specific to cassava breeding further exacerbated these effects in advanced cycles. Cassava reproduction is biologically constrained by low botanical seed production, irregular flowering, and variable seed viability. As a consequence, many crosses failed to generate sufficient progeny or produced seeds that were subsequently lost due to low germination rates and poor early seedling vigor. These factors substantially reduced the number of genotyped and evaluated individuals in later cycles, particularly C3, directly contributing to a reduction in population size and amplifying the effects of genetic drift. Both simulation-based and empirical studies consistently show that reduced effective population size not only increases inbreeding but can also impair the predictive efficiency of genomic selection models, especially in advanced cycles where genetic variance has already been eroded (Daetwyler et al., 2010; Gorjanc, Gaynor, and Hickey, 2018).

### Prospects for continued genetics gains in the cassava breeding pipelines

When considered jointly, the patterns observed for genetic diversity, predictive performance, and realized genetic gain provide important insights into the long-term sustainability of the breeding program. While genomic selection has clearly enhanced the efficiency and rate of genetic gain, the genetic shifts observed in C3 highlight the need for proactive strategies to mitigate the loss of genetic diversity and maintain effective population size. Among the most promising approaches is the deliberate expansion and renewal of the genetic base. Dos Santos et al. (2023a), for example, demonstrated the effectiveness of constructing core collections targeting key traits such as root quality, yield, and disease resistance. By capturing a reduced subset of individuals that retains a large proportion of the overall genetic diversity, such strategies can support the development of more representative and resilient training populations, as reflected in metrics including *H*_*o*_, *F*_*i*_, and overall genetic diversity.

Next, controlled recombination strategies across selection cycles offer a powerful means of balancing genetic gain and diversity. Approaches such as those proposed by Guimarães et al. (2025) explicitly guide crossing decisions to optimize long-term genetic variability while preserving favorable alleles. These strategies can be implemented through mating optimization frameworks, including mate selection (Wolfe et al., 2021) and optimum contribution selection (Meuwissen, 1997; Dagnachew and Meuwissen, 2016), which allow breeders to regulate parental contributions and control inbreeding without substantially compromising selection response.

Additional operational strategies may further enhance long-term sustainability, including the periodic introgression of novel parental lines, fine-tuning of selection intensity, and optimization of testing schemes through sparse or partially replicated trial designs. Their value can be further assessed using simulation platforms such as AlphaSimR (Gaynor, Gorjanc, and Hickey, 2021), which enable breeders to compare selection and crossing scenarios and evaluate expected impacts on genetic gain, inbreeding, and resource allocation across time horizons (Bančič et al., 2025).

Multivariate genomic selection also offers a promising way to improve predictive performance while moderating genetic divergence among traits. Because these models jointly predict correlated traits, they are particularly effective when traits show favorable genetic correlations and moderate to high heritability (Jia and Jannink, 2012; Okeke et al., 2017). By explicitly modeling genetic covariances, they not only enhance prediction accuracy but also clarify the structure of genetic variance, thereby supporting more balanced selection decisions when genomic estimated breeding values are incorporated into selection indices.

## Conclusions

The application of genomic selection resulted in significant genetic gains in industrially relevant traits in cassava, demonstrating its effectiveness as a core component of modern breeding pipelines. Although predictive ability varied modestly across selection cycles, it remained stable overall, and the consistent reduction in RMSE across generations indicates progressive improvements in the precision and calibration of genomic prediction models as the breeding population accumulated historical data.

Analyses of realized genetic gain revealed consistent responses for key productivity traits. Concurrently, targeted reductions in plant architecture scores reflected deliberate selection toward ideotypes better suited to mechanization, whereas modest declines in dry and starch content were attributable to indirect responses driven by underlying genetic correlations with yield-related traits. These outcomes highlight the inherent complexity of multi-trait improvement and reinforce the need for selection strategies that explicitly account for correlated responses, rather than focusing exclusively on single-trait gains.

Genetic diversity analyses further showed that recurrent genomic selection reshaped population structure across cycles, promoting genetic progress but also leading to a measurable reduction in genetic variability. These findings emphasize that sustained genetic gain under genomic selection requires not only accurate prediction but also the deliberate management of effective population size and recombination strategies to mitigate the erosion of diversity.

In general, this study underscores the central challenge faced by breeding programs that adopt high-accuracy genomic tools, balancing rapid genetic gain with the preservation of genetic diversity necessary for long-term progress and resilience. Periodic renewal or expansion of training populations, regulation of selection intensity, and the implementation of optimized mating and recombination schemes emerge as essential strategies to secure consistent short-term gains without compromising the long-term efficiency and sustainability of cassava breeding programs.

## Acknowledgment

The authors thank CNPq (Conselho Nacional de Desenvolvimento Cientıfico e Tecnológico), FAPESB (Fundação de Amparo à Pesquisa do Estado da Bahia), and CAPES (Coordenação de Aperfeiçoamento de Pessoal de Nıvel Superior) for financial support. This work was also supported by the NEXT-GEN Cassava project, through a grant to Cornell University by the Bill & Melinda Gates Foundation and the UK Department for International Development

This preprint was created using the LaPreprint template (https://github.com/roaldarbol/lapreprint) by Mikkel Roald-Arbøl .

